# Rare variant pathogenicity triage and inclusion of synonymous variants improves analysis of disease associations

**DOI:** 10.1101/272955

**Authors:** Ridge Dershem, Raghu P.R. Metpally, Kirk Jeffreys, Sarathbabu Krishnamurthy, Diane T. Smelser, David J. Carey, Michal Hershfinkel, Janet D. Robishaw, Gerda E. Breitwieser

## Abstract

Many G protein-coupled receptors (GPCRs) lack common variants that lead to reproducible genome-wide disease associations. Here we used rare variant approaches to assess the disease associations of 85 orphan or understudied GPCRs in an unselected cohort of 51,289 individuals. Rare loss-of-function variants, missense variants predicted to be pathogenic or likely pathogenic, and a subset of rare synonymous variants were used as independent data sets for sequence kernel association testing (SKAT). Strong, phenome-wide disease associations shared by two or more variant categories were found for 39% of the GPCRs. Validating the bioinformatics and SKAT analyses, functional characterization of rare missense and synonymous variants of GPR39, a Family A GPCR, showed altered expression and/or Zn2+-mediated signaling for members of both variant classes. Results support the utility of rare variant analyses for identifying disease associations for genes that lack common variants, while also highlighting the functional importance of rare synonymous variants.

**Author summary:** Rare variant approaches have emerged as a viable way to identify disease associations for genes without clinically important common variants. Rare synonymous variants are generally considered benign. We demonstrate that rare synonymous variants represent a potentially important dataset for deriving disease associations, here applied to analysis of a set of orphan or understudied GPCRs. Synonymous variants yielded disease associations in common with loss-of-function or missense variants in the same gene. We rationalize their associations with disease by confirming their impact on expression and agonist activation of a representative example, GPR39. This study highlights the importance of rare synonymous variants in human physiology, and argues for their routine inclusion in any comprehensive analysis of genomic variants as potential causes of disease.

## Introduction

The superfamily of G protein-coupled receptors (GPCRs) translates extracellular signals from light, metabolites and hormones into cellular changes, which makes them the targets of a significant fraction of drugs currently on the market [1]. Genome-wide association studies (GWAS) on common variants in GPCRs have begun to identify their contributions to various disease processes [2, 3]. However, many GPCRs lack common variants and alternate strategies are needed to understand their roles. Recently, sequence kernel association testing (SKAT) methods have been applied to rare variants in GPCRs to assess disease associations. Initial approaches binned all rare variants in a genomic region or gene [4, 5], but more recent methods group the rare variants most likely to contribute to disease associations, or aggregate variants based on domain or family structures [6, 7].

Large-scale studies of orphan or understudied GPCRs to characterize natural genetic variation in the human population can provide insights into the biological function and/or potential causal contributions to disease processes [8]. The DiscovEHR collaboration represents a tremendous resource [9]; here we use whole exome sequences and clinical information from 51,289 individuals in the DiscovEHR cohort. We identified sequence variants (common variants, mean allele frequency, MAF ≥ 1% and rare variants, MAF < 1%) for 85 orphan or understudied GPCRs not amenable to GWAS studies, and performed SKAT analyses. We binned rare variants into functional classes: loss-of-function (LOF, truncation and frameshift) variants; missense variants with predicted pathogenicity; or rare synonymous variants showing altered codon bias. We performed independent SKAT analyses on the various functional classes to determine their disease associations. Remarkably, for those GPCRs with sufficient numbers of patients for statistical analyses, we found the top disease associations were common to all available variant classes in the gene.

To assess the validity of the disease association results, we focused on *GPR39*. Among its particular advantages, *GPR39* contained sufficient numbers of variants in all three functional classes to be amenable to rare variant approaches. Second, *GPR39* is a small Family A GPCR, allowing rapid generation of mutants. Third, GPR39 is activated by extracellular Zn2+ and is coupled to inositol phosphate production [10, 11], permitting straightforward functional analyses. Finally, we have only a rudimentary understanding of the role(s) of GPR39 in human physiology, despite its broad expression and multiple cellular functions [12, 13]. Our results demonstrate the validity of combined computational and functional approaches to provide important insights into the clinical contributions of orphan and understudied GPCRs. Results also focus attention on the importance of rare synonymous variants for identifying disease associations. The overall strategy can be readily applied to other classes of genes without clinically important common variants.

## Results

### Rare variant-based disease associations of orphan / understudied GPCRs

We identified rare variants in whole exome sequences from 51,289 individuals, and to predict their functional impact, variants were annotated and sorted into three classes in order of predicted severity, i.e., Loss-of-Function (LOF; premature stop codon, loss of a start or stop codon, or disruption of a canonical splice site), missense variants (predicted to be likely pathogenic (pLP) or pathogenic (pP) by RMPath scoring (Supplemental Methods)), and synonymous variants with significantly different codon frequency relative to the reference codon (termed SYN_ΔCB, i.e., synonymous variants with altered codon bias).

Determining disease associations for genes having only low frequency variants requires binning of variants to increase statistical power. Binning methods have focused on gene or genetic region, and have recently been expanded to incorporate regulatory regions and/or pathways by incorporating biological information from curated knowledge databases [14]. For this study, we were specifically looking for clinically impactful rare coding variants which could be validated by functional studies of the relevant GPCRs. We used <u>s</u>equence <u>k</u>ernel <u>a</u>ssociation tests (SKAT) and binned variants according to functional classes described above, and focused on the disease associations which were common to more than one class. **Figure 1** illustrates the analysis pipeline for the entire set of 85 GPCRs. Of the 72 GPCRs with sufficient variants to allow at least 2 variant classes to be analyzed by SKAT, 28 yielded significant disease associations. Because this was an exploratory analysis, we used a significance cut-off of 0.05, i.e., P-values < 6.2E-07 after Bonferonni correction for multiple testing. **Table 1** shows the top disease associations for the 13 GPCRs that had sufficient variants in the LOF class. Supporting validity, associations between *GPR37L1* and epilepsy [15], and *LGR5* and alopecia [16, 17] have been previously identified by other means. Attesting to the discovery potential, other disease associations were novel, including *GPR161* and hyperhidrosis, *LGR4* and Sicca syndrome, *GPR153* and hyperpotassemia, *GPR84* and anemia, and *GPR39* and peripheral nerve damage, among others. Notably, the phenome-wide disease associations for the predicted LOF class showed congruence across the other variant classes for the subset of GPCRs having sufficient missense or SYN_ΔCB variants for independent analyses. While predicted LOF variants are easily identified and likely to have the greatest functional impact, missense variants classed as pLP + pP can also have significant effects on GPCR function. For the subset of GPCRs without sufficient LOF variants, we found significant disease associations for the missense and SYN_ΔCB classes of variants, **Table 1**. Some of the associations validate previous studies, including *GPR183* and disorders of liver [18], and *GPR85* with acute myocardial infarction [19]. Other associations were novel, and represent potential targets for future study, including *GPR132* and disturbances in sulphur-bearing amino acid metabolism, *GPR176* with asthma, *GPR12* with epilepsy, and *GPRC5D* with renal osteodystrophy. Of note, the SYN_ΔCB variant class had significant disease associations in common with LOF and/or missense variants, and most consistently had sufficient variants to be included in the analysis. Parallel SKAT analyses of the three distinct classes of rare variants produced two important results. First, independent analysis of distinct classes of variants yielded concordant disease associations, increasing confidence in the results. Second, the most consistent source of phenome-wide disease associations was SYN_ΔCB. Altogether, these results provide a strong rationale for including all three functional variant classes in disease association analyses.

**Table 1:**
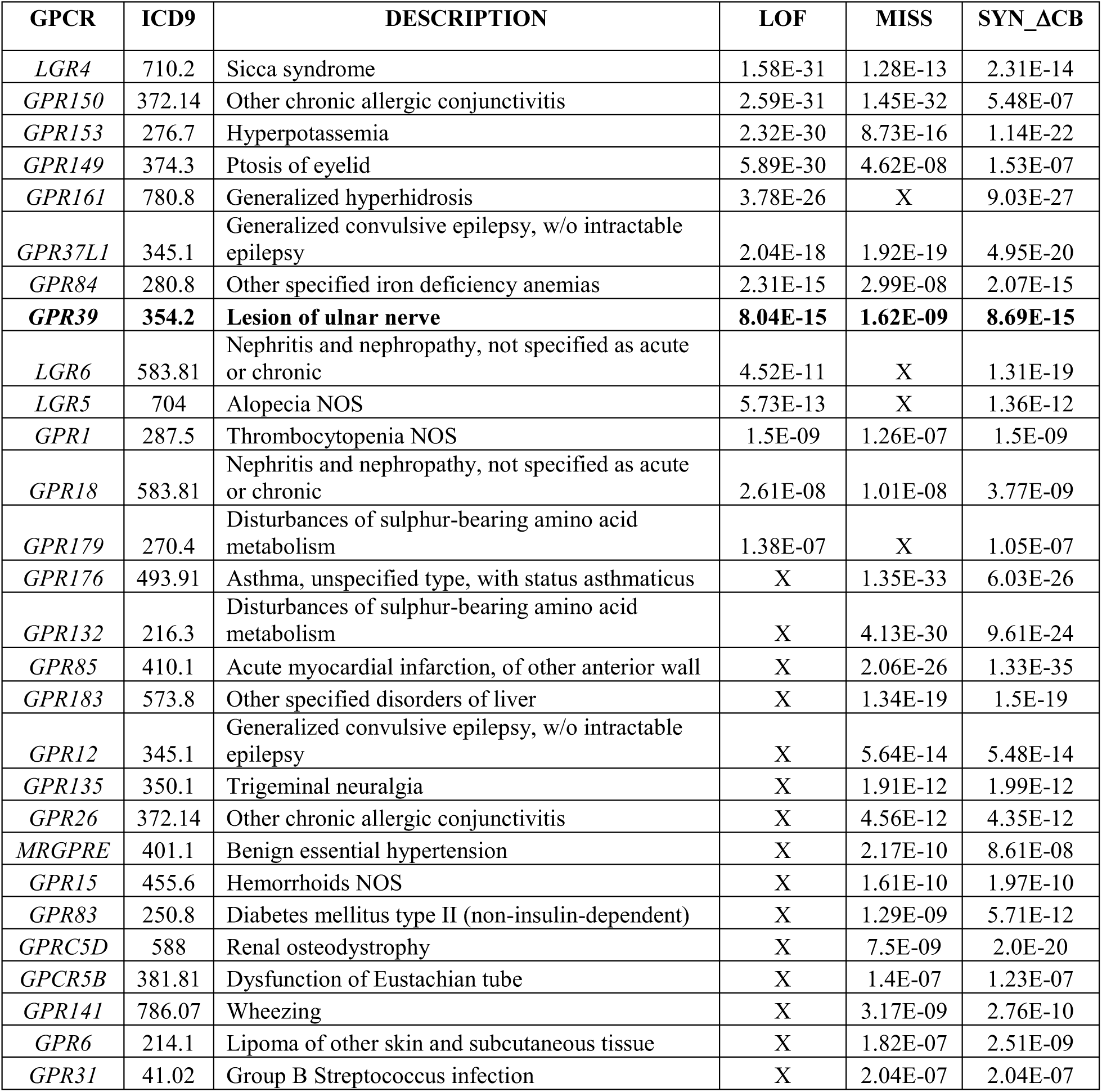
Orphan GPCRs with overlapping disease associations for multiple classes of rare variants. All orphan GPCRs with sufficient numbers of rare LOF, MISS or SYN_ΔCB variants in at least two categories were analyzed by SKAT. Absence of results in a category (marked by X) indicates insufficient number of variants and/or individuals having the variant for well-powered SKAT. A Bonferroni correction for multiple testing (123 sets, 656 ICD9 codes) was used to establish an exploratory significance cut-off of 0.05 (adjusted P-value threshold 6.2E-7). NOS, not otherwise specified.

**Figure 1:**
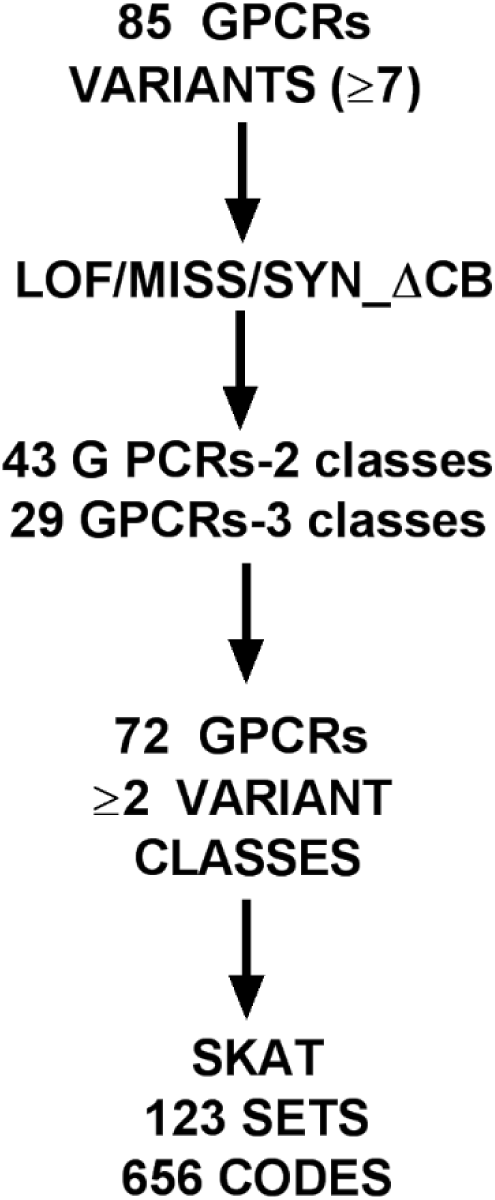
Analysis flow for orphan/understudied GPCR SKAT analysis. Rare variants from 85 GPCRs were identified in 51,289 WES dataset and sorted into LOF, missense and synonymous categories. Missense variants were further triaged to identify those predicted to be likely pathogenic or pathogenic by RMPath analysis (Supplemental Methods), and the subset of synonymous variants with large changes in codon usage bias were identified. Those GPCRs with at least 2 categories of variant having ≥ 7 distinct members were further analyzed (72 of 85 GPCRs). SKAT analysis was run independently on 123 sets of variants, against 656 disease codes having ≥ 200 individuals with ≥3 independent encounters in their EHR.

### Identification of *GPR39* common and rare variants in the 51,289 WES DiscovEHR cohort

Disease associations provide a rich source of hypotheses regarding the potential contributions of GPCRs. The first step in defining a causal role for these rare variants, however, is functional validation of the SKAT results. Here we focused in particular on rare synonymous variants with altered codon frequency (SYN_ΔCB), whose functional consequences are not well understood. Many of the GPCRs have no known agonist, and their dominant signaling pathways have not been characterized (**Table 1**). We therefore chose to focus on *GPR39*, the Zn2+ receptor, which has been characterized in cellular and knockout mouse models, and for which both agonist and dominant signaling pathways are known [12, 13].

All common and rare variants in *GPR39* were identified in 51,289 individuals in the DiscovEHR cohort [9] (**Figure 2)**. None of the *GPR39* variants were found in the ClinVar database. Accordingly, all LOF, missense, and synonymous variants were analyzed and classified as described for **Table 1**. The common variants were used for <u>P</u>henome-<u>w</u>ide <u>A</u>ssociation <u>S</u>tudies (PheWAS), and included a missense variant (2:133174764, Ala50Val) predicted to be benign, a synonymous variant (2:133174999; Thr128Thr), which changed a common to the rarest codon (acA (28%) to acG (12%), i.e., a 2.33-fold decrease in frequency), and a non-coding variant (2:133402607). Notably, no phenome-wide disease associations were detected (Supplemental **Figure S1**).

**Figure 2:**
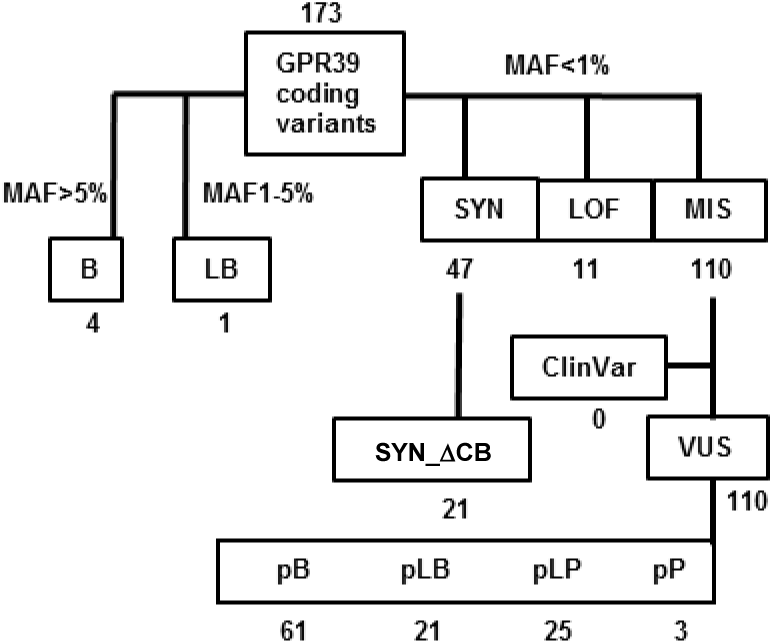
Pathogenicity triage of 159 coding variants in *GPR39* identified in 51,289 WES. Common variants with MAF≥1% were categorized as likely benign (MAF between 1-5%) and benign (MAF≥5%). All variants having MAF<1% sorted to synonymous, loss of function (LOF; frameshift or premature stop codon), and missense. Missense variants were thus Variants of Unknown Significance (VUS) and subjected to bioinformatics pathogenicity triage using RMPath pipeline, and classified as predicted benign, pB, likely benign, pLB, likely pathogenic, pLP or pathogenic pP.

The rare variants used for SKAT, i.e., LOF, missense (pLP + pP), and SYN_ΔCB variants, were distributed throughout the coding region of *GPR39* (**Figure 3**). Examination of the 39 missense variants that were categorized as pLP and pP confirmed that the RMPath pipeline identified variants which are likely to impact GPR39 function (**Figure 3**). The three pP variants (S78L, C108R, R133G) included a cysteine that participates in a disulfide bond at the extracellular face of GPR39 [20]. The pLP variants included one which contributes to the Zn2+ binding site (H19R) [21], two variants in a disulfide bond cysteine (C191G, C191W) [20], a residue in the highly conserved NPXXY motif common to Family A GPCRs (N340S) [22], and a variant distal to a putative palmitoylation site in the proximal carboxyl terminus (R352Q). Other residues in the pLP category introduce charges or proline residues into transmembrane helices or extracellular loops (S49N, M87I, T122M, A167S, P179T, R193C, S222Y, T291I, A293P, R302W, R302Q, A307V, P310A, Y314R, R320Y, S336R, R352Q, R362C, H367R, R386C, and R390C). We also identified the subset of 21 variants which introduced at least a two-fold change in site-specific codon frequency (SYN_ΔCB; denoted by arrowheads in **Figure 3**).

**Figure 3:**
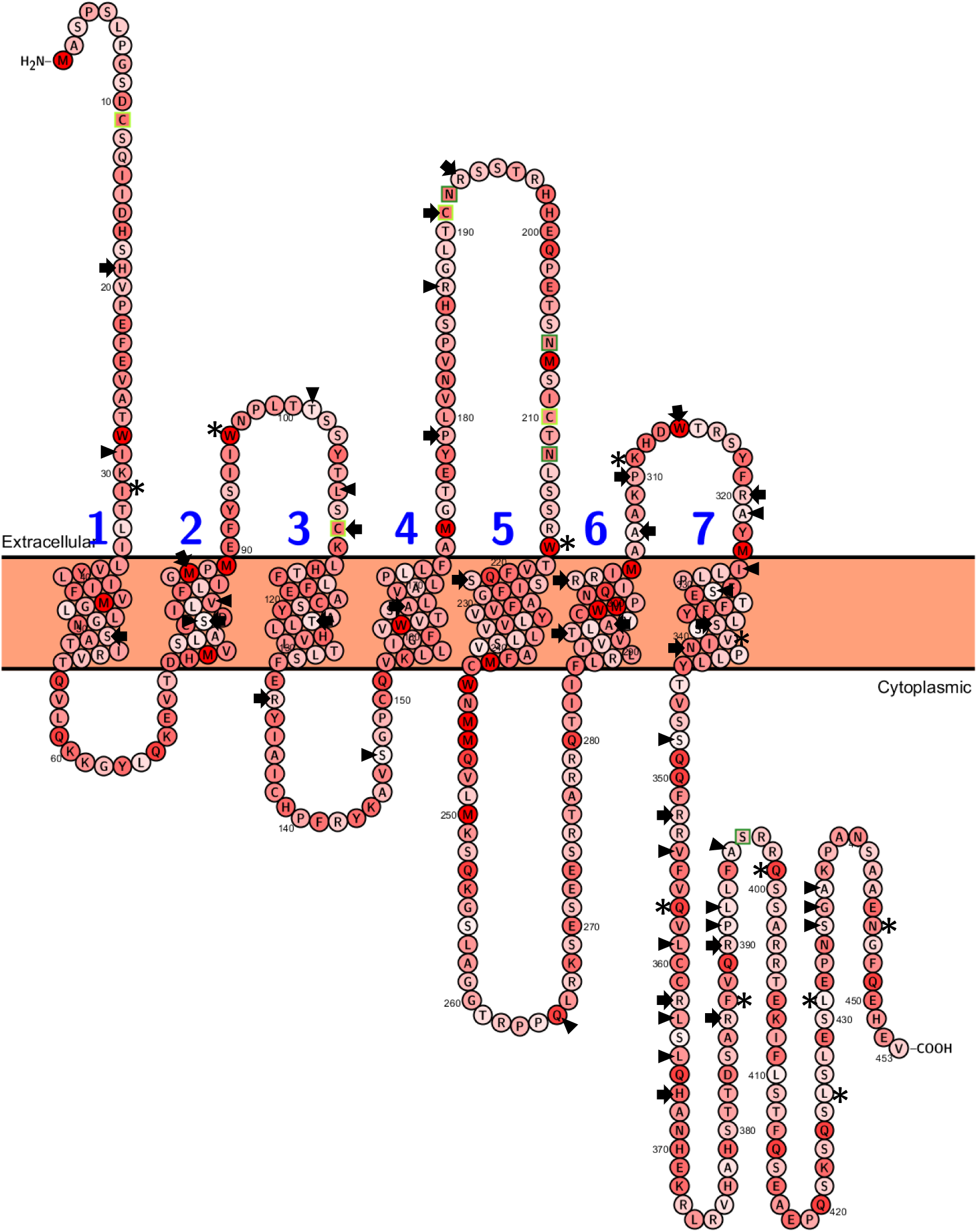
Snake plot of GPR39 topology indicating rare variants identified in 51,289 WES. The GPR39 sequence was color-coded to indicate reference codon usage frequency. For all amino acids having multiple potential codons, red indicates the most common codon, with progression from common to rare colored as gradations to light pink (rare). Arrowheads indicate the locations of rare synonymous variants exhibiting the largest changes in codon bias, either from common to rare or rare to common. Asterisks indicate rare LOF variants (truncations or frameshifts) and arrows indicate locations of rare missense variants combined for SKAT analysis.

### Disease associations of *GPR39* rare variant classes

The initial SKAT screen for *GPR39* disease associations compared top associations for the three independent classes of rare variants. To more fully explore the results for *GPR39* and increase power to identify all significant disease associations, we combined all individuals heterozygous for rare LOF, pLP and pP variants (termed LOF/MISS). Analysis was restricted to only those ICD9 codes with ≥200 individuals independent of genotype [23], and the results are plotted against ICD9 codes in **Figure 4A**. SKAT analysis was also run on rare SYN_ΔCB variants, plotted in **Figure 4B**. Both the LOF/MISS and SYN_ΔCB analyses had a significant number of phenome-wide associations in common. All results for both variant classes that were above the significance cut-off for multiple testing are listed in **Table 2**. Top associations for *GPR39* included lesion of the ulnar nerve and benign prostate hyperplasia with lower urinary tract obstruction. Note that individuals who had both rare LOF/MISS and SYN_ΔCB variants were excluded from analyses. Also of note in **Table 2**, some significant disease associations were specific to either the LOF/MISS or SYN_ΔCB variant classes. To assess the importance of rare variant pathogenicity triage for binning prior to SKAT analysis, we combined all rare variants (LOF, missense, and synonymous variants; 209 variants, 177 used in analysis) and ran SKAT. The only significant disease association was upper quadrant abdominal pain (ICD9 789.02; P-value 8.98E-06; significance cut-off at P < 7.6E-05). We also ran SKAT on those rare missense variants predicted to be benign (**Figure 2**, pB; 61 variants); no statistically significant disease associations were obtained. Likewise, we ran SKAT analysis on all rare synonymous variants and found top associations similar to those obtained for SYN_ΔCB, but with less significant P-values, suggesting dilution of impactful variants, i.e., lesion of ulnar nerve (P-value 1.08E-07), and benign prostate hyperplasia with urinary tract obstruction and other lower urinary symptoms (P-value 3.79E-05) (compare **Tables 2** and **3**). We conclude that pathogenicity triage to refine rare variant binning was crucial to identifying disease associations for the LOF and missense variants, and binning of the subset of rare synonymous variants having large changes in local codon frequency bias strengthened disease associations. Further, refining the rare variant input into the SKAT analyses yielded disease associations common to multiple independent classes of variants, strengthening confidence in the associations.

**Table 2:**
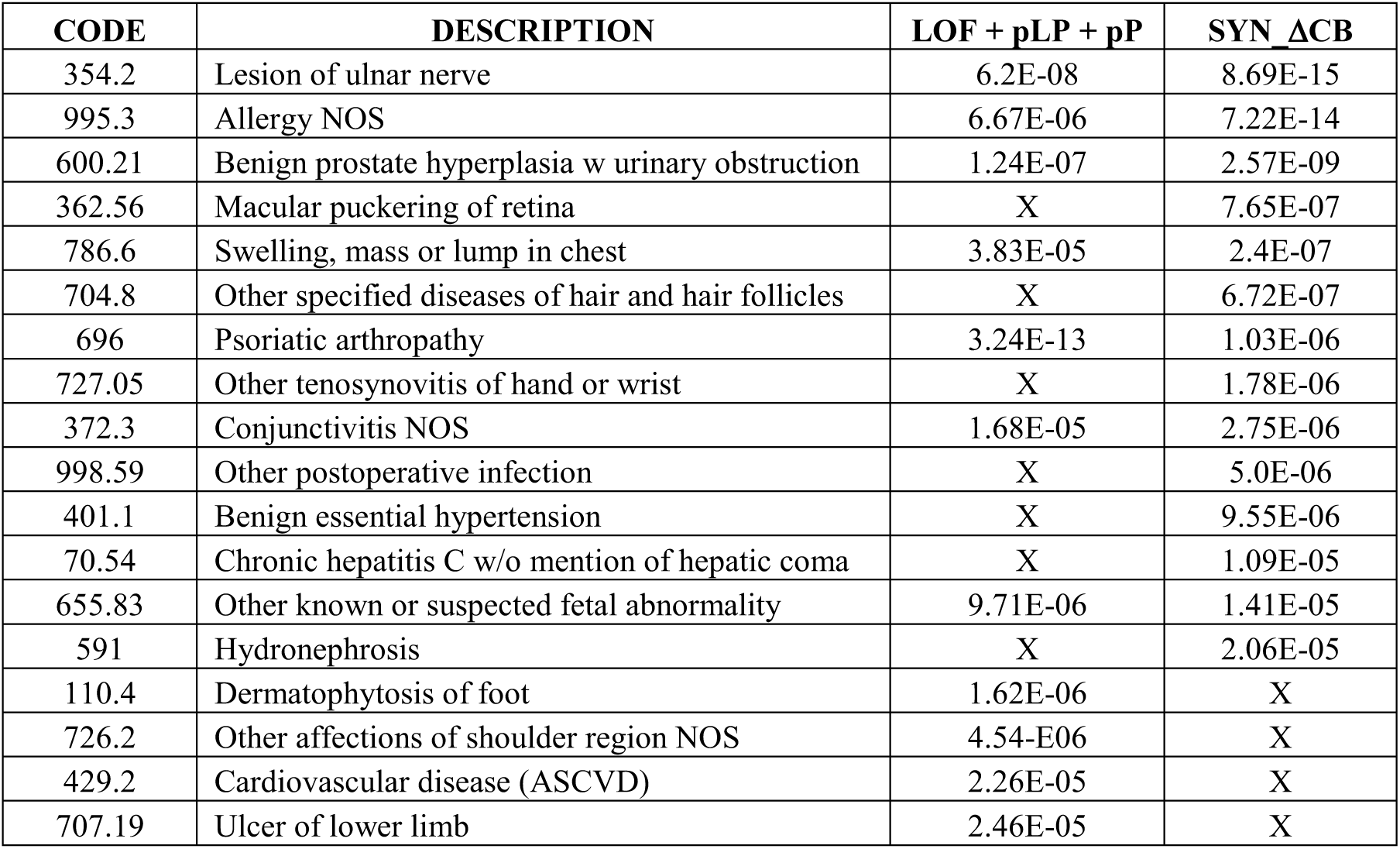
SKAT results for rare GPR39 variants in 51,289 DiscovEHR cohort. All LOF plus pathogenic missense (pLP + pP) were binned and analyzed by SKAT (53 markers; 221 individuals). The SYN_ΔCB were separately binned and analyzed (22 markers; 198 individuals). Listed are all significant associations when either/both variant categories had a P-value below the exploratory threshold of 0.05 (adjusted P-value threshold of 3.8E-5; 2 sets, 656 codes). Associations that were below the adjusted P-value threshold for only one variant category are also listed. NOS, not otherwise specified.

**Table 3:** SKAT analysis of all rare synonymous GPR39 variants in 51,289 DiscovEHR cohort. All rare synonymous variants were binned and analyzed by SKAT (46 markers; 940 individuals). Shown are all associations which were below the exploratory threshold of 0.05 (adjusted P-value 7.62E-05; 1 set, 656 codes). NOS, not otherwise specified.

| CODE | DESCRIPTION | P-value |
| --- | --- | --- |
| 696 | Psoriatic arthropathy | 3.87E-11 |
| 354.2 | Lesion of ulnar nerve | 1.08E-07 |
| 110.4 | Dermatophytosis of foot | 8.34-E07 |
| 726.2 | Other affectations of shoulder region, NOS | 2.24E-06 |
| 995.3 | Allergy, NOS | 3.51E-06 |
| 655.83 | Other known or suspected fetal abnormality | 4.38E-06 |
| 707.19 | Ulcer of lower limb | 1.06E-05 |
| 786.6 | Swelling, mass or lump in chest | 2.06E-05 |
| 429.2 | Cardiovascular disease (ASCVD) | 3.03E-05 |
| 649.13 | Obesity complicating pregnancy, childbirth, antepartum | 3.58E-05 |
| 600.21 | Benign prostate hyperplasia with lower urinary tract symptoms | 3.79E-05 |
| 440.9 | Atherosclerosis NOS | 6.31E-05 |
| 649 | Tobacco use disorder complicating pregnancy, childbirth | 6.8E-05 |
| 368.2 | Diplopia | 7.46E-05 |

**Figure 4:**
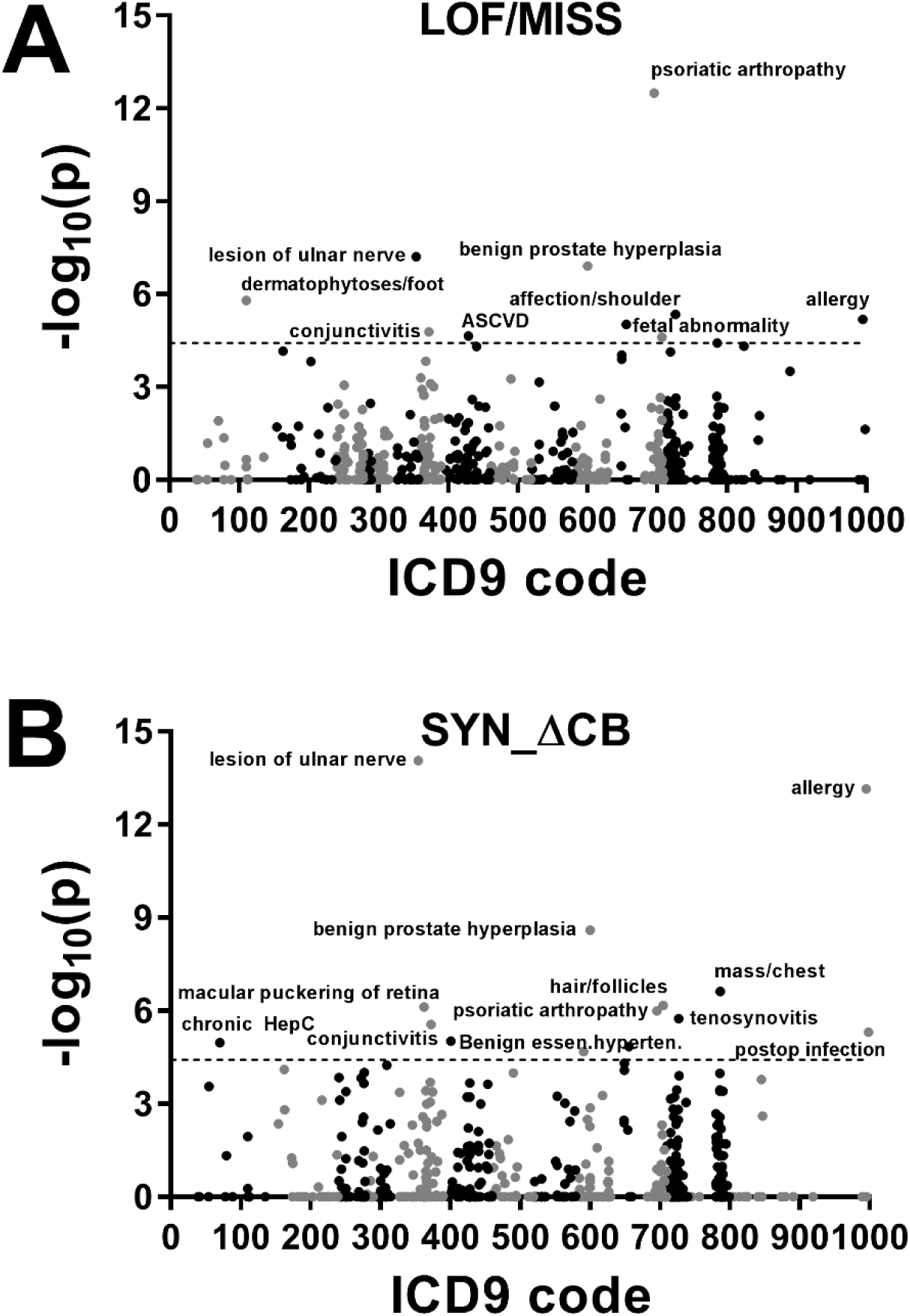
Manhattan plots of SKAT results for *GPR39* rare variants. A. Rare LOF and MISS variants were combined (39 total variants) and analysis run using SKAT. Top phenome-wide associations are labelled. B. Rare SYN_ΔCB variants (21 total variants) were analyzed with SKAT; top phenome-wide associations are labelled. Full details of associations common to both variant categories are presented in **Table 2**. For both **A.** & **B.**, the significance threshold was set at 0.05 (adjusted P-value 3.8E-05, 2 sets, 656 codes).

### Validation of functional impact of rare *GPR39* variants

To confirm the functional basis for the observed disease associations, we generated the rare missense and SYN_ΔCB variants in the reference human *GPR39* background. The cDNA construct included an amino terminal 3X FLAG epitope followed by a minimal bungarotoxin binding site, for ease of expression analysis and subcellular localization, respectively. Expression in HEK293 cells of equivalent amounts of cDNA and subsequent western blotting of lysates showed significant differences in net expression levels of the rare missense and SYN_ΔCB variants. Ten missense variants had significantly reduced expression and one had significantly increased expression relative to WT GPR39 (Supplemental **Figure S2**). Rare missense variants with altered expression were distributed throughout GPR39 structural domains, as color coded in **Figure 5A**. Likewise, SYN_ΔCB variant expression was significantly greater than WT GPR39 for three variants, while fourteen variants were expressed significantly less that WT (Supplemental **Figure S2**), quantified in **Figure 5B**. Surprisingly, despite the change from rare to common codon frequency for the majority of SYN_ΔCB variants, only four showed increased expression relative to WT. Four variants went from common to rare codons, denoted by * on the x-axis in **Figure 5B**. Contrary to expectations, Q265 was expressed at significantly higher levels than WT (∼150%), despite the large decrease in codon frequency. Overall, results argue that factors in addition to site-specific codon usage bias determine net expression levels for rare synonymous variants.

**Figure 5:**
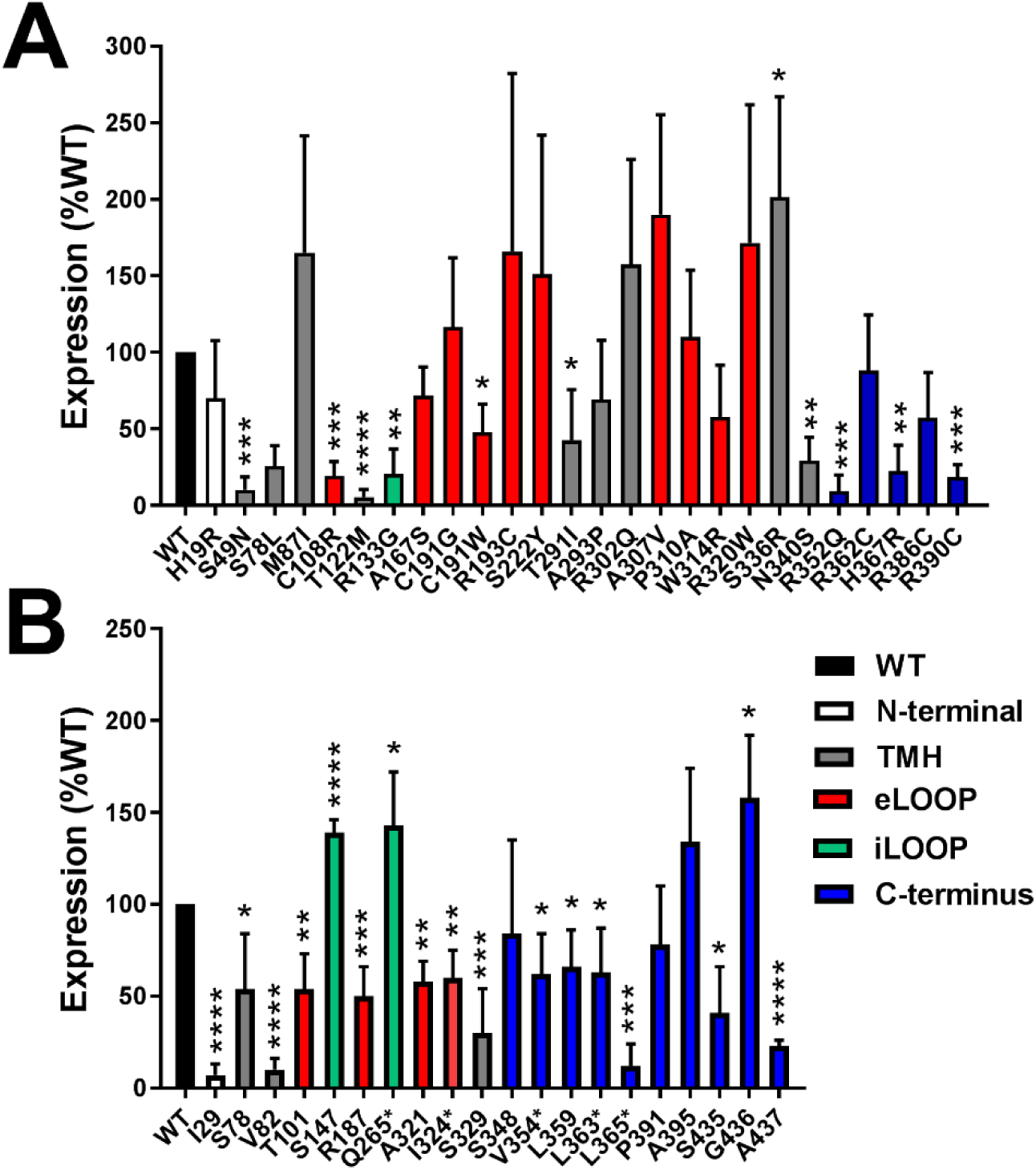
Expression of rare missense and synonymous GPR39 variants. A. Quantitation of expression levels of *GPR39* rare variants relative to WT. All blots had WT and HEK293 only lanes for background subtraction and normalization. Data presented as % of WT expression ±S.D. (n ≥ 3 independent transfections). Representative blots in **Figure S2**. **B.** Quantitation of expression levels of GPR39 rare SYN_ΔCB variants relative to WT. Details same as in **A**. Statistical analysis for both figures was two-tailed t-test relative to WT, with *p<0.05, ** p<0.01,***p<0.001 and ****p<0.0001.

GPR39 signaling is complex, with multiple signaling outputs. However, in both endogenous and heterologous expression systems, GPR39 activates inositol trisphosphate (IP3)-dependent release of intracellular Ca^2+^ [10, 11, 24]. The first step in the signaling pathway is Gq-mediated activation of phospholipase C, which produces diacylglycerol and IP3. Here we exposed HEK293 cells transiently expressing WT or variant GPR39 to its agonist, ZnCl2, and used a FRET assay to quantify accumulation of intracellular inositol monophosphate (IP1) in the presence of LiCl. **Figure 6A** illustrates normalized maximal levels of IP1 for WT and representative missense variants having decreased (R386C) or increased (W314R) EC50 for activation by ZnCl2. **Figure 6B** plots the EC50s for WT and all other missense variants, color-coded for their locations in GPR39 domains. Four variants, S78L, M87I, T122M and R133G, had no measurable activity, likely accounted for by low levels of expression (recall **Figure 5A**). Three additional variants, C108R, R193C and R302Q, had dose-response curves that were significantly right-shifted, precluding determination of EC50 or VMAX, despite expression levels not significantly different from WT. Likewise, **Figure 6C** illustrates the normalized dose-response relations for WT GPR39 and representative SYN_ΔCB variants which decrease (V354) or increase (I29) the EC50 for ZnCl2 activation, and **Figure 6D** plots the EC50s of WT GPR39 and all SYN_ΔCB variants. In contrast to the missense variants, all SYN_ΔCB variants had ZnCl2-stimulated IP1 activity. Six variants had increased EC50s and three variants having significantly reduced EC50s relative to WT. Surface expression of all rare missense and SYN_ΔCB variants was assessed by BgTx-Alexa594 binding on non-permeabilized cells (**Figures 6E, 6F**), and generally correlated with the extrapolated Vmax values for the IP1 assays (**Figures 6G, 6H**). Notable exceptions include the missense variants C108R, R193C and R320Q, which had surface expression comparable to WT but little IP1 activity. Likewise, the most striking SYN_ΔCB variant, Q265, had an overall increase in net expression (**Figure 5B**), increased EC50 relative to WT (**Figure 6D**), and increased surface localization and VMAX relative to WT (**Figure 6F, 6H**). Overall, results argue for a striking similarity in functional impact on expression and IP1 signaling for a subset of both rare missense and SYN_ΔCB variants of GPR39, rationalizing the disease associations identified for both classes of rare variants. The congruence of top disease associations between rare LOF, pathogenic missense and SYN_ΔCB variants for a significant subset of GPCRs (recall **Table 1**) argues strongly for the utility of these analytical approaches for identification of functionally impactful rare variants, and the imperative to include rare synonymous variants in genomic and functional analyses.

**Figure 6:**
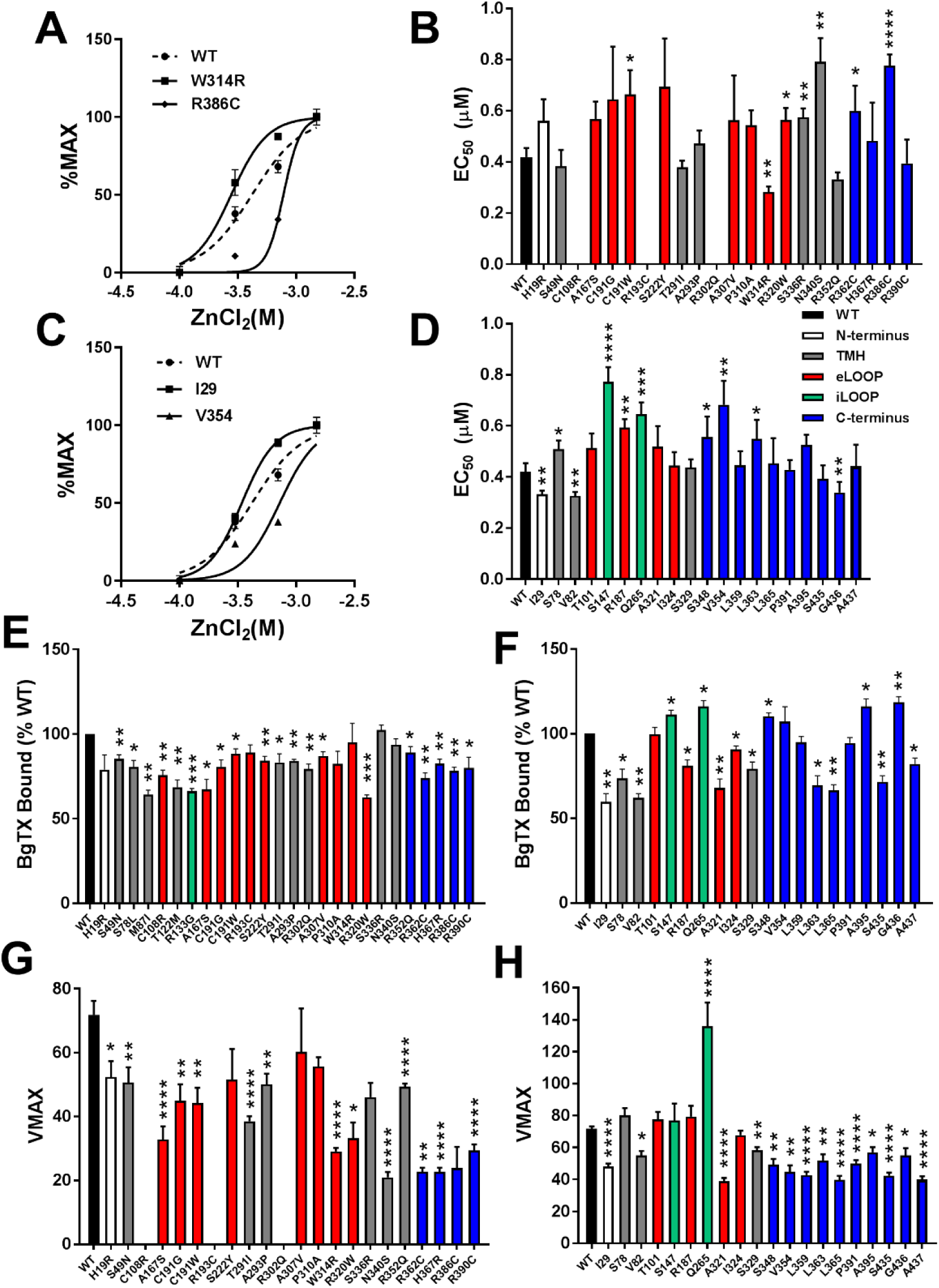
Functional analysis of rare missense and synonymous GPR39 variants. A. IP1 responses to various concentration of ZnCl2 for all variants that expressed ≥ 50% of WT. Representative curves and fits for WT, W314R and R386C are illustrated. Plotted are mean ± S.D. of three independent experiments, lines show fits to Michaelis-Menten equation. **B.** EC50s for all dose-response relations (as in **A**), plotted as mean ± S.E. for n=3 independent experiments. **C.** IP1 responses of representative SYN_ΔCB variants showing altered EC50s compared to WT, I29 and V354. Details as in **A**. **D.** EC50s for all dose-response relations (as in **A**) for SYN_ΔCB variants. **E.-F.** Bungarotoxin binding (see Methods) was used to assess surface localization of rare missense (**E**) and SYN_ΔCB (**F**) variants. **G.-H.** Extrapolated VMAXs of Michaelis-Menten fits to IP1 dose-response relations (see Methods). For all plots, *p<0.05, **p<0.01, ***p<0.001, and ****p<0.0001, by two-tailed t-test relative to WT, assuming unequal variances.

## Discussion

The related GWAS and PheWAS approaches, which utilize common variants, have produced strong, validated targets for drug development [2]. However, many genes do not have common variants, and additional approaches are required. The advantages of rare coding variant approaches are two-fold. First, rare variants can have stronger effects on protein function, and second, assessment of the impact of rare coding variants on protein expression and function is straightforward. However, due to their low frequency, it is necessary to combine rare coding variants to increase the statistical power for detecting disease associations. In this study, we used pathogenicity triage as a basis for rare variant binning, based on the hypothesis that aggregating variants with similar directions of effect on protein function would amplify disease associations. We considered, in order of the strength of their potential effects, predicted LOF variants, predicted likely pathogenic or pathogenic missense variants, and synonymous variants with altered codon frequency bias. as a test of the importance of rare variant triage on disease associations, we compared agnostic binning of all rare variants in *GPR39* with the disease associations identified for the three sorted variant classes. Of note, binning of all rare variants yielded no significant disease associations, while curated classes of those same rare variants had concordant top phenome-wide disease associations, obtained with non-overlapping individuals. Likewise, SKAT analysis of the subset of rare missense variants predicted to be benign yielded no significant disease associations. Results argue strongly that binning all rare variants in a gene or genetic region dilutes the effects of impactful variants, and appropriate pathogenicity triage of rare variants is an integral first step in successful application of disease association analyses.

Over the past 10 years, rare synonymous variants have been shown to impact protein expression and function with a frequency comparable to non-synonymous variants, although the mechanism(s) by which they exert their biological function are diverse [25–36]. Rare synonymous variants have not, however, been systematically utilized for disease association analysis [31, 37–39]. Here we demonstrate that 39% of understudied GPCRs have impactful rare synonymous variants which yield disease associations comparable to LOF or missense variants in the same gene, which may provide impetus for exploratory drug development despite the lack of identified endogenous agonists. Our data also highlight the necessity for improved approaches for pathogenicity triage of rare synonymous variants. For *GPR39*, SKAT analysis on all rare synonymous variants for the most part yielded disease associations similar to those obtained with the SYN_ΔCB subset, but having reduced significance, arguing for dilution of impactful variants. However, a few disease associations, including psoriatic arthropathy, showed more significant P-values in the combined synonymous variant analysis (3.87E-11) versus SYN_ΔCB (1.03E-06), implying that simply isolating those synonymous variants having altered local codon usage bias missed some variants with significant functional impacts (see **Tables 2** and **3**). Using both LOF/missense and synonymous variants as independent data sets in the SKAT analyses increases confidence in the identified disease associations and provided the impetus for further functional studies.

A disease association does not prove causation, but implies an impact on protein expression and/or function, which must be validated experimentally. As proof-of-concept, *GPR39* was chosen because its agonist and dominant signaling pathway were known, and rare variant classes produced strongly congruent disease associations. We identified 168 rare variants in *GPR39* in the 51,289 WES DiscovEHR cohort. Most individuals were heterozygous for the rare variants studied, and only 1/327 individuals had more than one rare variant. Of the 110 missense variants, the RMPath pipeline predicted that 28 were likely to be pathogenic (pLP + pP). Among these, rare variants previously shown to be important for GPR39 function were identified, including the zinc binding site [21], a critical extracellular disulfide bond [20], the pH sensor [11], and the NPXXY motif common to all Family A GPCRs [22]. When expressed in HEK293 cells, these variants had the expected effects on EC50s and maximal activities. Of the 47 synonymous variants, the codon usage table identified 21 variants having significantly altered codon usage frequency, and many had significant effects on expression, surface localization, and Zn^2+^-induced IP1 production. Surprisingly, the distributions of rare missense and SYN_ΔCB variants in the GPR39 structural domains were distinct. The majority of missense variants were localized to the transmembrane domain (77%), half in the extracellular loops (10/26), one an intracellular loop and the remainder in the transmembrane helices (9/26). In contrast, ∼50% of SYN_ΔCB variants were in the carboxyl terminus; the remainder were distributed among transmembrane helices (3/20), intracellular (2/20) and extracellular (4/20) loops. The carboxyl terminal variants in both classes had significant effects on expression and signaling, primarily reducing VMAX. The carboxyl terminus of GPR39 is relative large (108 amino acids) and has 22 putative phosphorylation sites. GPR39 signaling outputs are regulated by agonist-toggled binding of protein kinase inhibitor β [40]. In addition, phosphorylation by Rho kinase mediates desensitization of GPR39 [41]. Thus, carboxyl terminal missense or synonymous variants may alter secondary structure and/or phosphorylation to impact signaling. Results focus attention on the carboxyl terminus as a critical domain mediating distinct aspects of GPR39 signaling. It should be noted that GPR39 signaling is complex, including Gs-mediated cAMP production, Gq-mediated IP3 accumulation, and SRF-SRE-mediated transcriptional regulation via G12/13 [41]. The current study examined only Gq-mediated IP3 generation, leaving open the possible impact of missense and/or synonymous variants on signaling bias.

Based on ClinVar and related databases, *GPR39* has not been causally linked to disease. The present study focused on the general patient population, and all patients in the present study were heterozygous for *GPR39* rare variants. Nevertheless, both LOF/MISS and SYN_ΔCB variants yielded disease associations by SKAT which suggest a role for GPR39 in nerve function, benign hyperplasia of the prostate, psoriatic arthropathy, diseases of hair and hair follicles, and benign essential hypertension. These associations are in agreement with cellular and animal studies which suggest that GPR39 regulates neuronal transport of potassium and chloride, attenuating post-synaptic excitability [12, 42–44]. Animal models demonstrate GPR39-mediated up-regulation of the neuronal K+/Cl-cotransporter attenuates seizure activity [44], and knockout of GPR39 promotes depression [45]. GPR39 is up-regulated by some anti-depressants [46] in mouse models. One study in humans showed that GPR39 levels were significantly reduced in the hippocampus and cortex of suicide victims [47]. The highest levels of Zn2+ in the body are in the prostate and seminal fluid [48], and GPR39 is differentially expressed in normal versus malignant prostate cells, where it regulates cell growth and proliferation [10, 12, 13], potentially contributing to the association with benign prostate hyperplasia observed in the present work. Finally, the observation that individuals heterozygous for rare variants in GPR39 have a significant association with diseases of hair and hair follicles should be considered in light of a recent study that found that GPR39 marks a class of stem cells in the sebaceous gland and contributes to wound healing in mice [49]. Overall, the identification of a subset of clinical phenotypes that echo cell culture and animal model studies supports disparate roles for GPR39 in human physiology, and validates the SKAT approaches used herein to identify disease associations. Additional phenotypes, not previously associated with GPR39 including psoriatic arthropathy and macular puckering of the retina, deserve further attention. In conclusion, we have identified strong disease associations for GPR39 using either LOF/missense or rare synonymous variants, which confirm and extend cellular and animals studies. Rare variants from both the missense and synonymous classes altered GPR39 expression and function, providing a rationale for the observed disease associations. The approach applied to a larger set of orphan or understudied GPCRs also yielded robust associations with phenome-wide significance, providing potential novel targets for further investigation. The methods described in this report are easily applied to any set of genes of potential clinical importance with available genomic and integrated longitudinal electronic health records. Finally, this study highlights the importance of rare synonymous variants in human physiology, and argues for their routine inclusion in any comprehensive analysis of genomic variants as potential causes of disease.

## Materials and methods

### Whole Exome Sequence (WES) Data

The MyCode^®^ Community Health Initiative recruits Geisinger Health System (Geisinger) patients from a broad range of inpatient and outpatient clinics to integrate genetic information with their clinical EHR data to foster discoveries in health and disease [23]. For this study, we used WES from 51,289 participants [9], which includes patients enrolled through primary care and specialty outpatient clinics. Participants were 59% female, with a median age of 61 years, and predominantly Caucasian (98%) and non-Hispanic/Latino (95%). Patient demographics are in **Supplemental Table 1**. Whole exome sequencing was performed using 75 bp paired-end sequencing on an Illumina v4 HiSeq 2500 to a coverage depth sufficient to provide greater than 20x haploid read depth of over 85% of targeted bases for at least 96% of samples. Sample preparation, sequencing, sequence alignment, variant identification, genotype assignment and quality control steps were carried out as previously described [23].

### Analysis of site-specific rare codon changes

Codon usage tables define the relative fraction of different codons used in the genes of an organism [50, 51]. For all rare synonymous codons, we determined the ratio of the genome-wide codon fraction of the *GPR39* reference codon over the synonymous codon variant at that position, e.g., for Ser147Ser (tcG/tcA), 0.15/0.06 = 2.5, i.e., a 2.5-fold change. All synonymous variants with at least a two-fold change in site specific codon usage were used in the SKAT analyses.

### Phenome-wide association analysis (PheWAS)

De-identified EHR data was obtained from an approved data broker. All unique ICD9 codes with ≥200 patients (regardless of genotype) with at least 3 independent incidences of the ICD9 code were extracted from the EHR [52]. Individuals having 1-2 calls were excluded, and those having no calls of a particular ICD9 code were considered as controls. All non-Europeans were excluded from the analysis, as was one sample from pairs of closely related individuals up to first cousins. All models were adjusted for sex, age, age^2^ and first 4 principal components. The PheWAS R package were used for association analyses, which were performed for three GPR39 common variants (2:133174764, Ala50Val; 2:133174999; Thr128Thr; and 2:133402607, noncoding), and plotting of results was done using GraphPad Prism (V.6).

### Sequence Kernel Association Testing (SKAT)

To test for associations with clinical phenotypes, we used the sequence kernel association test (using default weights; SKAT R package), a gene-based binned statistical test comparing the burden of rare variants in cases and controls. Cases and controls were defined as for PheWAS analysis [52]. We defined various variant sets, including (1) only potential loss of function (splicing, stop gain, stop loss, or frameshift), (2) predicted pathogenic missense (pLP and pP by RMPath scoring) variants, and (3) synonymous variants having large codon usage frequency changes (at least a two-fold change in frequency between human reference codon and altered codon, SYN_ΔCB). For full *GPR39* analysis, we combined categories (1) and (2). Age, age^2^, sex, BMI, and first four principal components were used as co-variates. All non-European samples were excluded. Statistical significance was determined using the Bonferroni correction, α/m, where α=0.05 and m= # of variant sets and disease codes, as indicated for each analysis [53].

### Generation of *GPR39* variants

All missense and SYN_ΔCB variants were generated by gene synthesis (Thermo Fisher Scientific, Gene Art) based on the NP_001499.1 reference sequence (GeneID 2863, 453 amino acids), and included tandem tags at the amino terminus to facilitate expression and functional studies. After the initiation methionine codon, we added a 3X FLAG epitope followed by a minimal bungarotoxin binding site. The complete inserted sequence is: GACTACAAAGACCATGACGGTGATTATAAAGATCATGATATCGATTACAAGGATGACGATGACAAGGGTTGGAGATACTACGAGAGCTCCCTGGAGCCCTACCCTGACGGT.

### Expression of GPR39 variants

WT and variant constructs were transiently transfected into HEK293 cells. Cells were lysed after 48 hours, and 25 μg of lysates were separated on 4-15% SDS PAGE gels (BioRad), blotted to nitrocellulose, blocked with 5% milk in TBS-T, and exposed for 1 hour at room temperature to anti-FLAG monoclonal antibody conjugated to HRP (Sigma). Blots were exposed to SuperSignal West Pico Chemiluminescence Substrate, (Thermo Fisher), images recorded on a FUJIFILM LAS-4000mini luminescence analyzer and processed with Image-Gauge version 3.0. GPR39 expression was quantified using Multi-Gauge software, normalizing to WT expression and corrected for HEK293 background on the same blot.

### Functional analysis of GPR39 variants

WT and variant constructs were transiently transfected into HEK293 cells, cultured for 24 hours, and equivalent numbers of cells were then plated in poly-D-lysine-coated 96 well plates for a further 24 hours. Replicates were treated with various concentrations of ZnCl2 for 10 min at 37°C. Levels of IP1 were determined as recommended by manufacturer (IP-One ELISA kit, Cisbio USA, Inc.). Plates were read on a POLARStar Omega (BMG Labtech) plate reader. EC50 was determined by fitting normalized dose-response data with the Michaelis-Menten equation, and VMAX was separately calculated on raw data (GraphPad Prism, V6). Parallel plates were exposed to Alexa-594-conjugated bungarotoxin (3 μg/ml, 5 min at 37 °C, 5% CO2) to assess surface expression of GPR39 variants. After removal of bungarotoxin and washes with PBS, fluorescence was read on the POLARStar Omega (BMG Labtech) reader. Each variant was assayed in ≥3 independent transfections; WT, standards and solution blanks were included in each assay.

## Acknowledgements

We are grateful to Geisinger for financial support, Geisinger and the Regeneron Genetics Center for access to the DiscovEHR cohort, and to the patients in the MyCode Biobank.

## Supplemental Methods

**In-silico <u>R</u>adical <u>M</u>utations <u>P</u>athogenicity (RMPath) prediction pipeline.**

Rare variants are categorized into two groups: (1) nonsense and frame-shift variants, and (2) missense variants, and were scored by summing numerical values derived from predictions.

**(1) Scoring of nonsense and frame-shift variants.**

SnpEff fields: LOF[*].PERC ≥ 0.5 or NMD[*].NUMTR > 3 are considered potential Loss of Function (LOF) variants.

**(2)Scoring of missense variants.**

1. SIFT scoring: D (damaging) = 2; All others = 0.
2. PolyPhen2 scoring: (i) HDIV: Probably damaging (D) = 2; Possibly damaging (P) = 1; all other results = 0. (ii) HVAR: Probably damaging (D) = 2; Possibly damaging (P) = 1; all other results = 0.
3. MutationTaster scoring: Disease causing (D) = 2; all other results = 0.
4. MutationAssessor scoring: High (H) = 2; Medium (M) = 1; all other results = 0.
5. FATHMM scoring: D (damaging) = 2; all other results = 0.
6. LRT: D(deleterious) → score 2; All others → score 0.
7. PROVEAN scoring: D(damaging) = 2; all other results = 0.
8. GRANTHAM scoring: ≥140 = 4; ≥120 = 3; ≥90 = 2; ≥70 = 1; all other results = 0.
9. Genomic Evolutionary Rate Profiling GERP++ rejected substitutions (RS) scoring: ≥6 = 4; ≥5= 3; ≥3.5 = 2; ≥2 = 1; all other results = 0.

Scores from all tools were summed to generate a cumulative score (= RMPath score), with a maximum possible value of 24. Variants with RMPath scores ≤8 were predicted to be Benign (pB), those with scores in the range from 6 to 11 were predicted to be Likely Benign (pLB); those with scores between 11 and 17 were predicted to be Likely Pathogenic (pLP); and those with scores ≥17 were predicted to be Pathogenic (pP). For clinical purposes, all variants are considered Variants of Unknown Significance (VUS) until family co-segregation and/or functional analyses definitively establish a variant as pathogenic.

**Supplemental Table 1.** Demographics of the DiscovEHR cohort used in the present study. Abbreviations, EHR, electronic health record, BMI, body mass index.

| Basic demographics | DiscovEHR Sequenced Patients |
| --- | --- |
| Total patients (N) | 51,289 |
| Female, N (%) | 30,290 (59%) |
| Median age, yrs | 61 (range 48-72) |
| Median BMI, kg/m<sup>2</sup> | 30 (range 26-36) |
| Median years of EHR data | 14 (range 11-17) |
| Median medication orders/patient | 129 (range 37-221) |

**Supplemental Figure S1.**
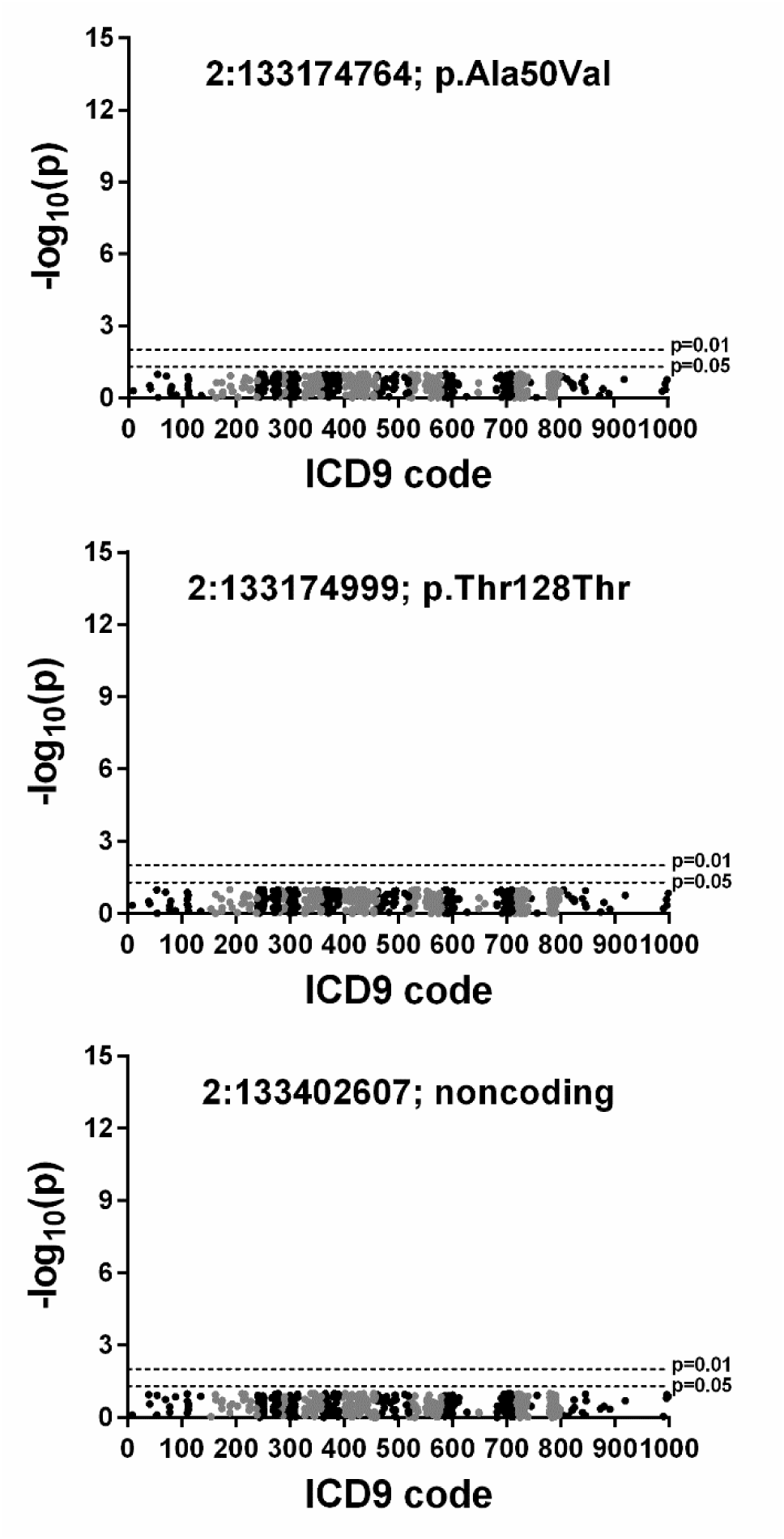
Manhattan plots of results of PheWAS analyses of common GPR39 variants. No association results exceeded the minimal P-value threshold of 0.05, uncorrected for multiple testing.

**Supplemental Figure S2.**
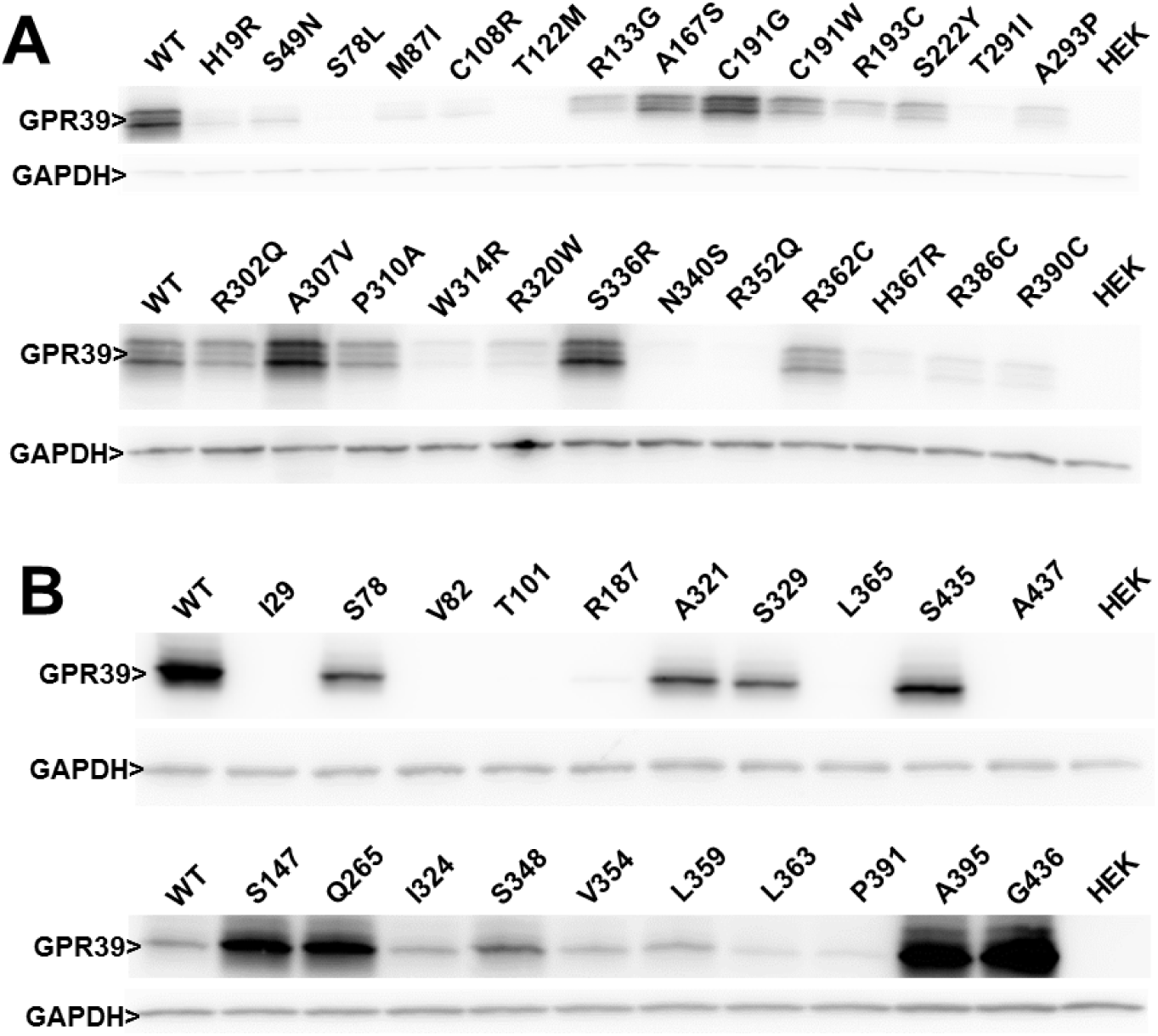
Representative western blots of rare missense and synonymous variants identified in the 51,289 WES cohort. Western blots of HEK293 cells lysates expressing WT or rare variant missense or SYN_ΔCB cDNAs were run as described in Methods. **A.** Missense variants run in order from amino to carboxyl terminal location; each gel also had WT and untransfected HEK samples for normalization; GAPDH was used as loading control. **B.** Expression of rare synonymous variants was more variable, and rare variants were sorted for high expression (top row) or low expression (bottom row) for better quantitation; WT and HEK were run on each gel for normalization; GAPDH was used as loading control.

